# Aurora kinase A enables collective invasion and metastasis by endowing a leader cell phenotype and stabilizing cohesion with follower cells

**DOI:** 10.64898/2026.03.31.715024

**Authors:** Bo Peng Zhou, Tony L.H. Chu, Aidan Gallant, Ryan Ghorayeb, Shanshan Wang, Tariq A. Bhat, Chelsey Gough, Roderic Espin Garcia, Miguel Angel Pujana, C. James Lim, Christopher A. Maxwell

## Abstract

The metastatic process initiates with collective cell invasion into surrounding tissues and axillary nodes, and subsequent colonization at a distant site. Previously, we found collective invasion is augmented during the G2 cell cycle phase, facilitated through Aurora kinase A (AURKA)-mediated centrosome polarization in the leader cell. Here, we identify cell cycle-associated gene signatures as overrepresented in axilla and liver metastatic sites, with AURKA expression strongly correlated with breast cancer metastasis signatures, and pan-cancer patient survival. Then, we show GFP-AURKA expression endows breast epithelia cells with the ability to form outgrowths within immune-incompetent chicken embryos. Multi-parametric imaging of wound closure assays reveals phenotypes enabled by, and dependent upon AURKA expression. We discover leader cells express AURKA and acquire front-polarized centrosomes, which differentiates them from other cells in the migrating group. Ectopic expression of GFP-AURKA induces a leader cell phenotype. Conversely, inhibition of AURKA activity alters actin dynamics, promotes turnover of cell contacts, and reduces coordination within migrating groups. Specifically, AURKA interacts with the actin regulator EPLIN, and AURKA inhibition localizes EPLIN to lamellipodia and away from E-cadherin-positive contacts. Inhibiting these necessary roles for AURKA may provide a critical barrier against the metastatic spread of human breast carcinoma cells.

## Introduction

Breast cancer is the second leading cause of cancer-associated death in women. Breast cancer survival is significantly reduced following the transition of the tumor from localized to invasive and metastatic disease [1]. The metastatic process, in turn, is believed to initiate with breast carcinoma cell invasion into surrounding tissues and axillary nodes, and subsequent colonization at a distant site, such as the liver [2]. Carcinoma cells that guide collective invasion often acquire distinctive molecular programs compared to the tumor cells at the original site [3], and invasion is frequently accompanied by so-called leader cells guiding finger-like cellular protrusions [4]. Leader cells provide directionality cues and are linked to the migrating group by E-cadherin complexes and actomyosin cables, which restrict the protrusion of adjacent cells at the wound edge [5, 6]. Leader cells also display other defining characteristics: they extend large, actin-rich lamellipodia, exhibit front-rear polarity, and acquire distinctive molecular features [3, 7], such as the expression of cytokeratins, like Keratin 14, and filaments associated with epithelial (E) to mesenchymal (M) transition (EMT), like vimentin [3, 8].

Single-cell resolution and spatial expression analysis of the metastatic breast tissue ecosystem, including stromal, myeloid, lymphoid, and malignant tissues, indicates strong inter-patient variability of expression profiles, but spatial organization of malignant cells with partitioning of EMT phenotypes, which are characterized by distinct cell cycle genes [9]. The relationship between cell cycle progression, EMT expression profiles, and migratory behaviour is likely to be complex and dynamic; indeed, single-cell, spatially-resolved transcriptome analysis of human breast cancers revealed negative correlation between EMT-enriched and proliferation-enriched or cell cycle-enriched states across all cancer epithelia regions [10]. Likewise, invasive regions within primary tumors, experimental models of metastasis, and circulating tumor cells each display heterogeneous, hybrid E/M areas, with E-cadherin+/ Vimentin+ regions [8], as well as dynamic transitions [6, 11]. Previously, we reported that carcinoma cells acquire some features of leader cells in culture, as they naturally progress through the cell cycle, including front-rear polarity and expression of vimentin, and these transitions rely on Aurora kinase A (AURKA) activity at the centrosome [12].

The expression of *AURKA*, along with the expression of *estrogen receptor* (*ESR1*) and *human epidermal growth factor receptor 2* (*ERRB2*), categorizes major breast cancer subtypes. For example, low *AURKA* expression is indicative of Luminal A breast cancer with favorable survival [13]. Conversely, elevated expression of *AURKA* associates with poor patient survival [14], younger age and aggressive features [15], EMT [16], and chemoresistance [17]; indeed, the *AURKA* loci at 20q13 is a recurrent amplification event in breast cancer metastasis [18]. Mechanistically, AURKA promotes centrosome maturation and the formation of a bipolar spindle during mitosis [19], but it also regulates cytoskeleton dynamics in non-mitotic cells [20]. In this study, we show that ectopic GFP-AURKA expression endows a non-cancer breast cell-line with the ability to form outgrowths within immune-incompetent chicken embryos. AURKA expression fosters the emergence of leader cells and promotes the cohesion of follower cells through the maintenance of cell contacts. Specifically, inhibition of AURKA activity mislocalizes the actin regulator EPLIN from E-cadherin-positive cell contacts to lamellipodia at the leading edge. Together, these data demonstrate that AURKA expression is sufficient to enable collective invasion and metastasis making its inhibition a potential therapeutic strategy to prevent metastasis of human breast carcinoma cells.

## Materials and Methods

### Ethical regulations

The studies using the chicken embryo chorioallantoic membrane (CAM) model followed BCCHR institution internal protocols and University of British Columbia animal committee guidelines. Experiments with the CAM model adhered to Canadian Council of Animal Care (CCAC) regulations that did not require approval and ARRIVE guidelines 2.0 [21].

### Analysis of TCGA gene-expression and clinical data

TCGA gene-expression and clinical data [22] were analyzed as previously described [23]. Single-sample gene-set enrichment scores were calculated from primary-tumor RNA-seq data using ssGSEA implemented in the GSVA R package [24] (version 1.43.1). Pearson correlations between *AURKA* expression and 6,356 curated MSigDB gene-set [25] scores were calculated across all tumors and within PAM50 molecular subtypes. *P* values were adjusted for multiple testing using the Benjamini–Hochberg approach. Gene sets annotated with “metastasis” or “invasion” and showing a FDR-adjusted *P* < 0.05 in at least one group were selected for visualization. Association between *AURKA* tumor expression and disease outcome were assessed by Kaplan–Meier analysis after stratifying samples into expression tertiles or into low/high groups relative to the cohort mean; differences between survival curves were evaluated using log-rank tests. For pan-cancer analyses, TCGA overall-survival and progression-free-interval data were analyzed using Cox proportional-hazards regression, with *AURKA* expression as a continuous variable and adjustment for age, gender, and tumor stage when available. Hazard ratios and 95% confidence intervals for *AURKA* were estimated separately for each cancer type.

### Cell culture

MCF10A RFP-TUBA1B were purchased from Sigma-Aldrich (CLL1039). Parental MCF10A cells were obtained from Dr. J Brugge (Harvard University, Boston MA). All cell lines were grown at 37 °C in a 5% (v/v) CO2 incubator. All cell lines were periodically checked for mycoplasma using a mycoplasma detection kit (Abcam, Vancouver, Canada).

MCF10A sublines were cultured in Brugge media, which is DMEM/F12 (1:1) media supplemented with 5% horse serum, 20 ng/mL epidermal growth factor (EGF), 10 µg/mL human insulin, 0.5 µg/mL hydrocortisone, and cholera toxin 100 ng/mL. MCF7 and MDA-MB-231 cells were grown in DMEM (high glucose) supplemented with 10% FBS and P/S.

### Virus packaging and transduction

The pHR_dSV40-Aurora A-GFP plasmid (Addgene, #67924) and the pHR_dSV40-GFP plasmid (a gift from Dr. James Lim) were transfected and packaged in lentivirus using psPAX2 and pmD2.G plasmids, which were gifts from Didier Trono (Addgene plasmids #12260 and 12259). Production and collection of lentiviral particles were performed as described by the CalPho Mammalian Transfection Kit (Clontech).

MCF10A RFP-TUBA1B cells were transduced with lentiviral particles containing either GFP-alone (control) or GFP-AURKA constructs. 72 hours post lentiviral transduction, GFP-positive cells were isolated using flow cytometry, cloned, and three clones of each treatment were selected for further analysis.

To generate cell lines stably expressing the luciferase reporter gene, parental MCF10A RFP-TUBA1B cells and cells stably expressing GFP-AURKA were infected with lentivirus that encodes for the click beetle luciferase-GFP protein (transfer plasmid is pELNS.CBG-T2A-GFP). 72 hours after lentiviral transduction, cells were subjected to flow cytometry to isolate cells that were GFP-positive. Cells co-expressing GFP-AURKA and luciferase-GFP were cloned, and three clones were selected for further analysis. Stable transfection was verified by routinely checking the cells for fluorescence by microscopy throughout the duration of the investigation. We validated luciferase expression for each cell line using bioluminescence imaging, as well as immunofluorescence and western blot analysis.

### Introduction of human MCF10A cells into chick chorioallantoic membrane (CAM)

Fertilized eggs from White Leghorn chickens (*Gallus gallus domesticus*) were purchased from the University of Alberta, Edmonton, Alberta. Following overnight reset of the air sac at 14°C, eggs were placed into a hatcher incubator (Digital 1502 sportsman, GQF, Berryhill, ON Canada) at 37°C with >70% humidity. On embryonic day 4 (E4), eggs were cracked with a rotary cutting tool (Dremel) and put in lidded sterile weigh boats (Cat. No. 08-732-113, Fisher Scientific). The ex ovo prepared CAMs were further incubated until E13.

Before each CAM experiment, luciferase expression was confirmed in the cells. For onplant implantation, MCF10A cells (1 x 10^6^ per onplant) were harvested and resuspended 1:1 in media and Geltrex (Lot#A1413.02 & A14132-02, Gibco, NY USA). 40 µL droplets were plated onto pre-warmed dishes using the hanging drop method. After setting at 37°C for 30 minutes, the onplants were incubated in complete media for 24 hours. They were then transferred onto gently lacerated E13 CAMs, imaged daily, and harvested on day 5 or 6 for further analysis. To assess the invasive and metastatic potential, GFP-luc and GFP-AURKA/GFP-luc cells (1 x 10^6^ per embryo) were resuspended in 50 µL of Dulbecco’s phosphate-buffered saline (DPBS, Gibco, Waltham, MA) and kept on ice until intra-amniotic (IA) injection into E13 embryos. Following injection, embryos were monitored daily for viability for 6-8 days prior to bioluminescence imaging for luciferase activity.

### Bioluminescence imaging and analysis

All bioluminescence imaging was performed on the Ami-X in vivo imaging system (Spectral Instruments Imaging, Tucson, AZ). At the study endpoint, embryos received 100uL of D-luciferin (GoldBio) via the amniotic cavity and were incubated for 5-15 minutes. Images were acquired in vivo and post-mortem for organ-specific invasion and metastasis analysis with the following settings: <30s exposure, binning of medium-high, anti-glare selected, 1.2 Fstop, and open emission (no filter). Data were plotted and statistical analysis was performed on GraphPad Prism. Total flux (photons/seconds) was calculated by subtracting the background signal from the embyro’s region of interest (ROI). Following image acquisition, tissues were harvested and formalin-fixed for paraffin embedding (FFPE) and sectioning.

### Histology and FFPE immunofluorescence

After extraction, tissues were fixed in 10% neutral buffered formalin overnight at room temperature before being embedded in paraffin and sectioned at a width of 5 microns. H&E-stained tissue sections were prepared and imaged to identify malignancies (MAPcore, Vancouver, Canada).

FFPE samples were kept at 67°C for 20 mins before being washed repeatedly in a sequential manner, with xylene, 100% EtOH, 95% EtOH, 70% EtOH, 50% EtOH and water. Then, the samples were fully submerged in citrate antigen retrieval buffer (pH 6) and boiled for 30 minutes in a rice cooker. The samples were allowed to cool to room temperature before proceeding. Each section was then blocked with 1% BSA 5% horse serum in PBS for 1 h at room temperature, permeabilized with 0.3% Triton-X 100 in PBS, then incubated with primary antibody at 4°C overnight, secondary antibody at room temperature for 2 hours, and used the Vector TrueVIEW Autofluorescence Quenching Kit (Vector Laboratories, SP-8400-15) per the manufacturer’s recommendation before incubating with Hoechst 33342 (1: 2 000) for 10 minutes at room temperature and mounting with VECTASHIELD Vibrance Antifade Mounting Medium. All slides were stored at 4°C until analysis.

### Wound closure assay

Cells were grown in standard growth media until 90% confluence. Cells were then serum-starved for 24 hours to promote G1-synchronization [12] and ensure minimal proliferation during the first 6 hours of wound closure. Wounds were made using a P200 pipette tip. Plates were then washed to remove cell debris before changing back to standard growth media and cells were allowed to migrate.

For wound closure assays using MLN8237 (1 μM, 2.5 μM, 5 μM) or MG132 (1 μM), after making the wound with a P200 pipette tip, each well was washed with PBS before adding standard growth media with DMSO. Cells were allowed to migrate for 3 or 6 hours before either changing to media with or without MLN8237 or MG132.

Cells were imaged using an ImageXpress Micro High Content Screening System (Molecular Devices, Inc) or Incucyte® S3 Live-Cell Analysis System (Sartorius, Inc). Image analysis was performed using MetaXpress and ImageJ.

### Immunofluorescence

All cell lines were seeded onto autoclaved coverslips and grown until the desired confluency. Then, the media was removed and coverslips were washed three times with PBS. The cells were fixed with either 4% paraformaldehyde in PBS for 15 min at room temperature or ice-cold methanol for 3 min at -20°C. Then, the cells were blocked with 3% BSA/PBS for 1 hr at room temperature, then incubated with primary antibody for 1 hour at room temperature or overnight at 4°C, incubated with secondary antibody, incubated with Hoechst 33342 (1: 5 000) for 10 minutes at room temperature, and then mounted with Prolong Gold Antifade Mounting Reagent (Life Technologies, P36935) and stored at 4°C until analysis. Images acquired with a confocal laser-scanning microscope (Fluoview Fv10i, Olympus; Leica SP8, Leica; CellDiscovrer7, Zeiss).

The following primary antibodies were used: Alexa-647-beta-tubulin (TUBB) (ThermoFisher, MA5-16308-A647, 1:100), K14 (Abcam,181595, 1:1 000), cytokeratin 14 (Abcam, ab181595, 1:1000), GFP (Abcam, ab183734, 1:200), phosphorylated AURKA (pAURKA) (Cell Signalling, 3079, 1:1 000), AURKA (Abcam, ab13824, 1:200 or Cell Signaling, 3092S, 1:400), E-cadherin (Cell Signaling, 3195S, 1:300 or Cell Signaling, 14472S, 1:300), EPLIN (Cell Signalling, 16639-1-AP, 1:100 or Cell Signalling, sc-136399, 1:100) and gamma-tubulin (TUBG1) (Sigma, T6557, 1:1 000). Antibodies conjugated with Alexa-488, Alexa-549, and Alexa-647 (Life Technologies) were used as secondary antibodies.

Image analysis was performed using ImageJ. To measure E-cadherin intensity at cell-cell junctions, the line scan function in ImageJ was used and drawn from the center of the leader cell nucleus to the nearest follower cell nucleus and the midpoint of the line was centered at the cell-cell junction.

### Immunoprecipitation and western blot analysis

Cells were lysed with chilled NP40 immunoprecipitation buffer (50 mM Tris-HCL, pH 7.4, 150 mM NaCl, cOmplete™, Mini, EDTA-free Protease Inhibitor Cocktail, PhosSTOP™, 1% NP40) on a rotator in the cold room. Lysates were clarified by centrifugation and incubated with Dynabeads Protein A (Invitrogen, 10002D) overnight then incubated for 2 h at 4°C with the following antibodies: EPLIN (NB100-2310, Novus Biologicals), AURKA (3092S, Cell Signaling) or Rabbit IgG (AB-105-C, RD Systems). Beads were washed with washing buffer (50 mM Tris-HCL, pH 7.4, 150 mM NaCl, cOmplete™, Mini, EDTA-free Protease Inhibitor Cocktail, PhosSTOP™, 0.1% NP40) and then eluted. Samples were separated by SDS-PAGE and analysed by western blot.

### Live-cell imaging of the cytoskeleton

To visualize cytoskeletal dynamics, SPY555-tubulin (Cytoskeleton, CY-SC203) and SPY650-FastAct™ (Cytoskeleton, CY-SC505) was diluted 1000x into complete MCF10A media without phenol red and then added to MCF10A RFP-TUBA1B cells 2 hours post-scratch. Live-cell imaging of cell migration was captured using CellDiscoverer7 (Zeiss) with 20x objective or 50x water-immersion, at an interval of 10-20 minutes.

Images were processed by making a leader cell mask using the tubulin and actin dyes to only crop the leader cell from the rest of migrating cells. Then, the nucleus centroid was used to center the cell before making inverted images. Temporal-color coded hyperstack of leader cells showing overlay of the time-lapse of tubulin and actin were made using non-centered, cropped leader cells. ImageJ was used to process these images.

### Live-cell imaging of lysosomes

Cells were grown on collagen coated glass bottom plates and a wound closure assay was performed. Cells were stained with LysoTracker™ Deep Red (ThermoFisher, L12492, 75nM) and SPY555-tubulin for 2 hours before media change and imaging. Confocal immunofluorescence imaging (Cell Discoverer 7, Zeiss) was used to capture migrating cells. Whole cell masks were drawn around leader cells using phase contrast. Then, automatic thresholding of the tubulin channel was used to define centrosome regions (ZEN).

### Analysis of published single cell RNA sequencing dataset

Data for the single cell RNA sequencing analysis was downloaded as a processed Seurat object from cellxgene, which included cell type annotations, G2/M scores, and normalized gene expression values used for all analyses. For pathway enrichment, Hallmark Gene sets were directly loaded into RStudio using the msigdbr package, and single cell pathway analysis [26] was performed.

### Statistical methods

Statistical analysis was performed using GraphPad Prism v5.01 for Windows (Graphpad Software). Pairwise comparisons were made using unpaired Student’s t-test. Comparisons of multiple groups were made using One-way ANOVA with a Bonferroni post-test.

## Results

### AURKA expression is augmented at metastatic sites and correlates with molecular signatures for metastasis and pan-cancer survival

Single-cell RNA sequencing was recently used to profile fresh tumor biopsies from a cohort of 30 patients with metastatic breast cancer, including malignant cells captured from liver (66, 293 cells from n=16 patients), axilla (5 765 cells from n=3 patients), and breast (2 283 cells from n=2 patients) sites of disease [9]. To mine this dataset for insight into the metastatic process, we performed single cell pathway analysis [26] on the annotated cells (**Fig. 1a**) [9]. We focused on hallmark gene sets annotated in the Molecular Signatures Database (MSigDB) [25] and analyzed differential gene expression within malignant cells in “axilla versus breast” or “liver versus breast”. This analysis revealed a significant enrichment of “G2M checkpoint” and “mitotic spindle” pathways in both the axilla sites and the liver sites relative to the primary breast tumor site (**Fig. 1b**) (**Supplementary Fig. 1a**). Conversely, the EMT pathway was enriched at the primary breast tumor site (**Fig. 1b**) (**Supplementary Fig. 1a**), which is consistent with a prior report of negative correlation between EMT-enriched and proliferation/cell cycle-enriched states [10]. We found significant overlap between high G2/M scores in this dataset and *AURKA* expression (**Fig. 1a**). To further evaluate the association between *AURKA* expression and cancer metastasis or invasion, we performed a meta-analysis of the >6 000 curated and annotated gene sets in the MSigDB [25]. For each gene set or pathway annotated with the term “metastasis” or “invasion” where UP denote genes that promotes cancer metastasis and DOWN (DN) the contrary, we computed a score and found significant positive correlation between these scores and the expression of AURKA across tumors and breast cancer subtypes. Almost all gene sets annotated with UP in metastasis or invasion are strongly and positively correlated with AURKA expression, while DN shows the opposite (**Fig. 1c**).

**Figure 1.**
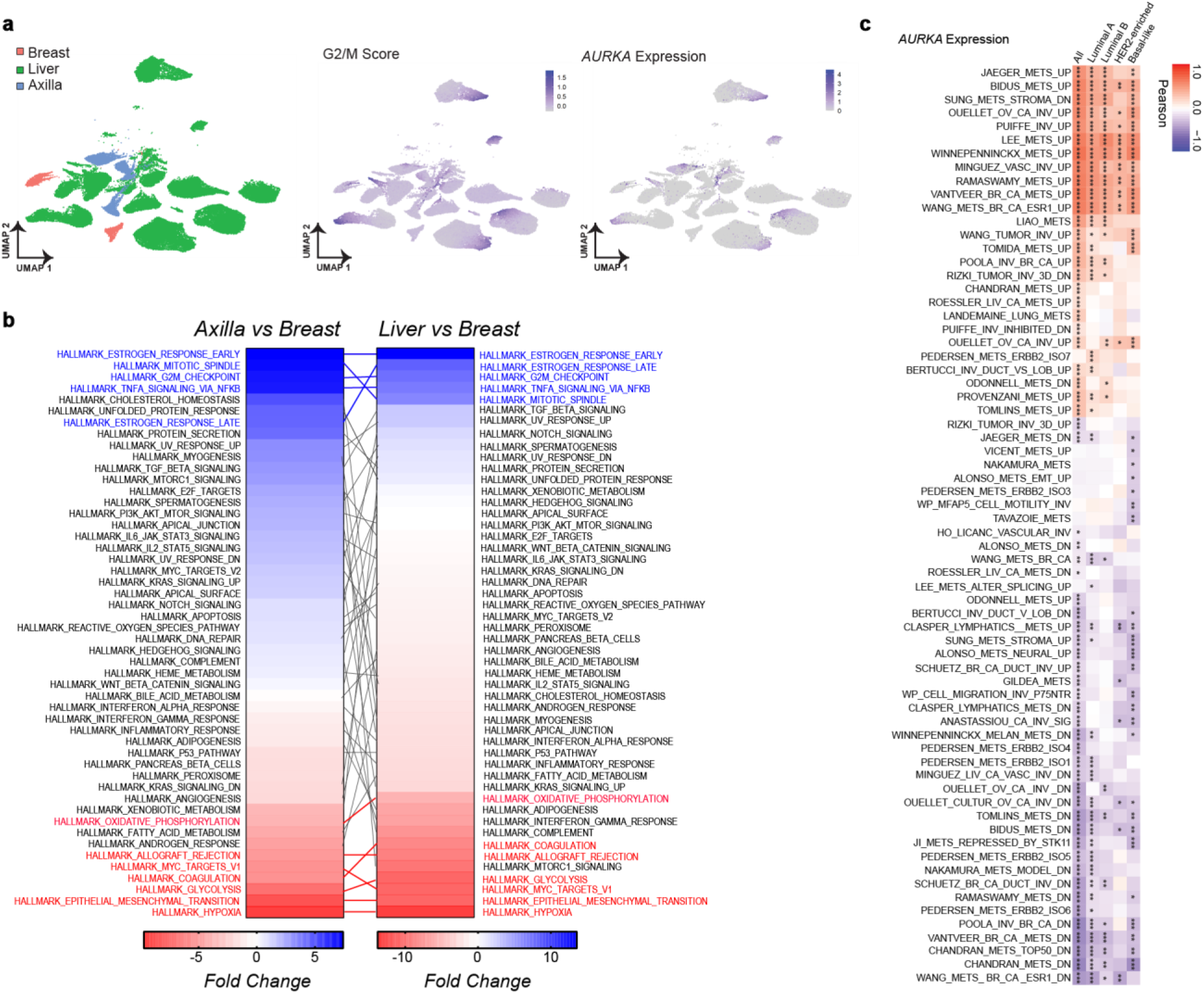
*AURKA* expression is augmented at metastatic sites and correlates with molecular signatures for metastasis and pan-cancer survival. **a.** Uniform manifold approximation and projection (UMAP) including all malignant cells from breast, liver, and axilla biopsies. G2/M scores and *AURKA* expression are overlayed onto the UMAP. Data sourced from [9]. **b.** Hallmark Gene Sets plot of fold change differences for Axilla versus Breast and Liver versus Breast highlighting gene sets that are both significantly different and have an absolute fold change value of 5 or more are highlighted with sets enriched in breast (red) or enriched at metastatic sites (blue). Data sourced from [9]. **c.** Heatmap of correlation between *AURKA* expression against gene sets/pathways annotated with “metastasis” or “invasion” in Molecular Signatures Database of GSEA. * indicate FDR-adjusted p-value. Data sourced from [25].

AURKA expression predicts breast cancer survival [14]. Indeed, our analysis of the METABRIC dataset revealed a decrease in the median overall survival (OS) and disease-free survival (DFS) of all breast cancer patients with high *AURKA* expression compared to patients with low *AURKA* expression (**Supplementary Fig. 1b**). To examine the broader relationship between *AURKA* expression and pan-cancer survival, we calculated the hazard ratio (HR) for OS and progression-free interval (PFI) of *AURKA* expression across TCGA studies (**Supplementary Fig. 1c**). Notably, of the 18 cancer types examined, we observe statistically significant increased HR with *AURKA* expression for OS of breast cancer (BRCA, HR=1.17, p=0.018), head and neck squamous cell carcinoma (HNSC, HR=1.3, p=0.0085), kidney renal cell clear cancer (KIRC, HR=1.47, p=5.85×10-6), liver hepatocellular carcinoma (LIHC, HR=1.27, p=0.0045), lung adenocarcinoma (LUAD, HR=1.26, p=0.00074), pancreatic adenocarcinoma (PAAD, HR=1.49, p=0.00074), prostate adenocarcinoma (PRAD, HR=2.16, p=0.023) and uterine corpus endometrial carcinoma (UCEC, HR=1.43, p=0.0019) (**Supplementary Fig. 1c**). Moreover, we found similar significantly increased HR with *AURKA* expression for PFI across multiple cancer sites (**Supplementary Fig. 1c**). Overall, we conclude that *AURKA* expression has a significant negative association with OS and PFI across cancer sites, and a strong positive association with breast cancer metastasis or invasion signatures, suggesting a capacity to promote cancer metastasis and invasion.

### GFP-AURKA expression enhances outgrowth potential in a chick embryo model

We evaluated the capacity for elevated AURKA expression to promote outgrowths of non-malignant, human breast cells using MCF10A RFP-TUBA1B cells, which have red fluorescence protein (RFP) attached to the endogenous *tubulin alpha 1B* (*TUBA1B*) locus. First, we introduced GFP-control (GFP-CTRL) or GFP-AURKA into MCF10A RFP-TUBA1B parental cells generating three clones for each and confirming their expression by western blot analysis (**Supplementary Fig. 2a**). Next, we introduced GFP-luciferase into MCF10A RFP-TUBA1B cells, named GFP-luc, or into GFP-AURKA clone #3, named GFP-AURKA/GFP-luc #3, and confirmed GFP-AURKA expression through western blot analysis (**Fig. 2a**) (**Supplementary Fig. 2b**) and immunofluorescence analysis (**Supplementary Fig. 3a,b**). Then, using cell number titration and bioluminescence imaging, we confirmed luciferase expression was equivalent across GFP-luc and GFP-AURKA/GFP-luc #3 cells (**Supplementary Fig. 3c**).

**Figure 2.**
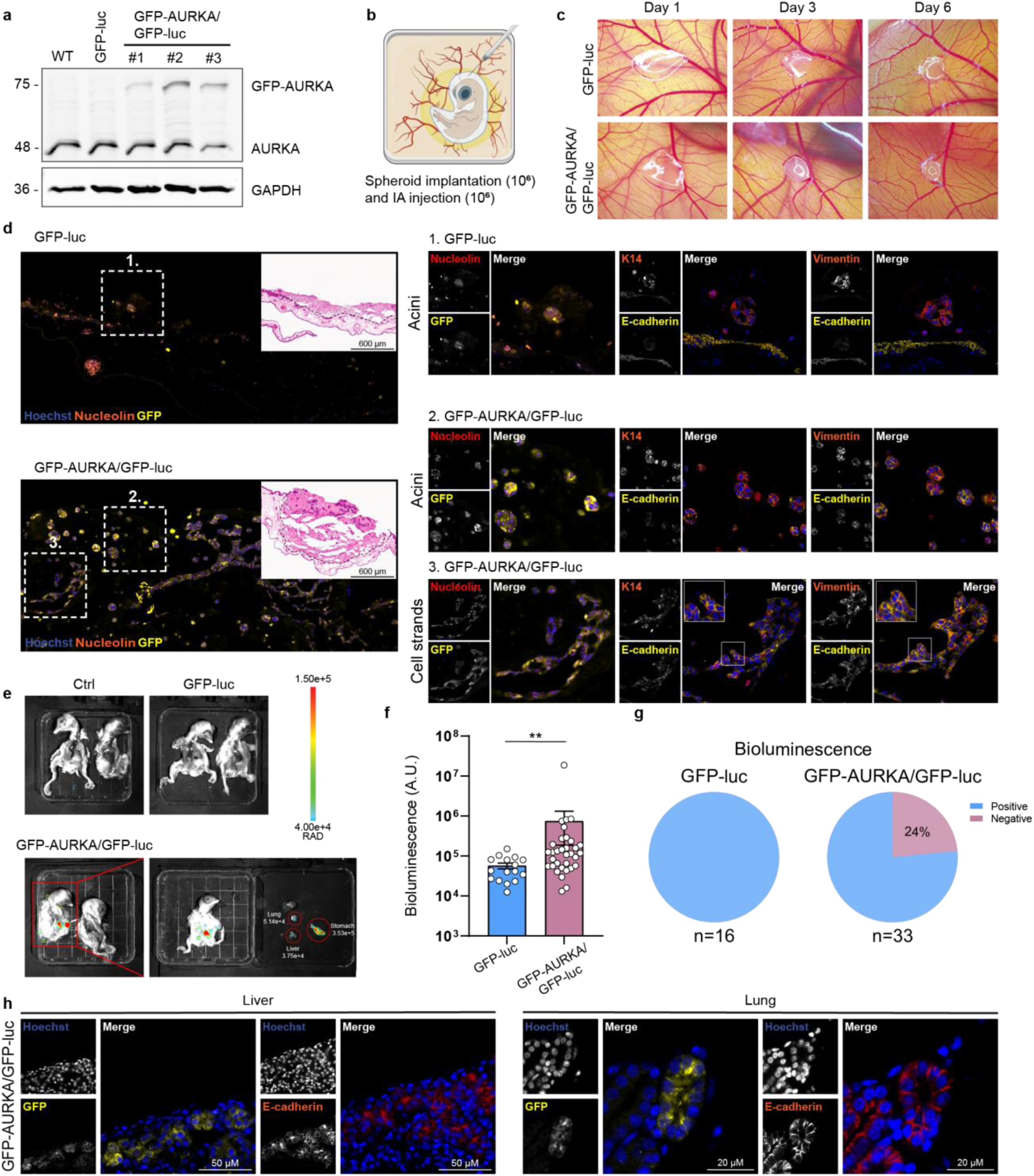
GFP-AURKA expression enhances metastatic potential in a chicken embryo model. **a.** Western blot analysis of parental MCF10A RFP-TUBA1B (WT), GFP-luc and GFP-AURKA/GFP-luc clones, stained for AURKA and GAPDH. **b.** MCF10A cells were either implanted onto the CAM as a onplant or injected into the amniotic cavity. **c.** Representative brightfield images over 6 days of growth for onplants (1 x 10^6^ cells per onplant) following their engraftment onto embryonic day (ED) 13 chick embryos. **d.** H&E images (inset) and immunofluorescence (IF) analysis of FFPE CAM isolated at day 6. The presence of human cells was confirmed through IF detection of GFP and a human-specific nucleolin. Human cells were also stained for their expression of K14 and E-cadherin, or vimentin and E-cadherin. Human cells were found as acini structures in both (1) GFP-luc and (2) GFP-AURKA/GFP-luc injected onplants. Human cells were found as cell strand structures in (3) GFP-AURKA/GFP-luc injected onplants. The tips of cell strand structures (boxes) showed positive staining for K14/E-cadherin/vimentin suggestive of a hybrid EMT phenotype. **e.** Representative bioluminescence imaging, through IVIS, of chick embryos collected six days after injection with either GFP-luc or GFP-AURKA/GFP-luc cells. **f.** Quantitation of background subtracted whole chicken embryo bioluminescence imaging of embryos injected with either GFP-luc cells or GFP-AURKA/GFP-luc cells. P values calculated by using the Mann Whitney U-test. **g.** Summary of injection experiments for GFP-luc cells (n=16) or GFP-AURKA/GFP-luc (n=33). **h.** Immunofluorescence analysis of luciferase-positive chick embryo tissues, previously injected with GFP-AURKA/GFP-luc cells, stained with GFP or E-cadherin.

To assess the ability of MCF10A RFP-TUBA1B clones to engraft *in vivo*, one million cells were embedded in Geltrex, cultured as onplants for 24 hours, and then implanted onto a lacerated chicken embryo CAM (**Fig. 2b**). Brightfield images were taken of the onplants over the course of 6 days of incubation (**Fig. 2c**). On day 1, the spreading pattern of GFP-luc and GFP-AURKA/GFP-luc onplants on the CAM surface was similar however, by day 3, GFP-AURKA/GFP-luc onplants were distinctive with a denser core persisting to day 6 of engraftment (**Fig. 2c**), which may be indicative of improved engraftment [27].

Onplants were isolated at day 7 following engraftment, fixed and then embedded in paraffin for subsequent H&E staining and immunofluorescence detection of GFP-positive human cells as well as epithelia (E-cadherin) and migratory markers (K14, vimentin). The boundary between the chicken embryo CAM and the Geltrex-embedded onplants were visualized in the H&E stained tissues (inset, **Fig. 2d**) (**Supplementary Figure 4a**) enabling the analysis of MCF10A cell engraftment, cell survival and growth. Human MCF10A cells were identified through immunofluorescence detection of GFP as well as human-specific nucleolin. Because chicken red blood cells are nucleated, their potential autofluorescence was quenched (**Supplementary Figure 4b,c**). In H&E stained tissues of onplants containing either of GFP-luc or GFP-luc/GFP AURKA MCF10A RFP-TUBA1B cells, acini-like structures were present after six days of engraftment; immunofluorescence detection of GFP and human-specific nucleolin confirmed these structures to be comprised of human cells (**Fig. 2d**) suggesting the acinus structures formed in Geltrex during the 24 hours prior to engraftment and are sustained within the chicken embryo CAM. In addition, GFP-AURKA/GFP-luc onplants showed more extensive growth, including the presence of cellular strands that resemble invasive cellular collectives (**Fig. 2d**) (**Supplementary Fig. 4d**). For this reason, we examined the architecture of these cell clusters through the detection of K14, E-cadherin and vimentin, and found the GFP-AURKA/GFP-luc strands to have heterogeneous expression of all three markers (**Fig. 2d**). Therefore, GFP-AURKA expression is sufficient to enhance the engraftment potential and/or survival of non-cancer MCF10A RFP-TUBA1B breast epithelial cells.

To examine whether GFP-AURKA expression is sufficient to increase the outgrowth and spread of non-cancer MCF10A RFP-TUBA1B cells, we injected cells into the intra-amniotic cavity of the chicken embryo, and measured bioluminescence with IVIS after 1 week (**Fig. 2e,f**) (**Supplementary Figure 5a**). At endpoint following intra-amniotic injection, for a subset of injected embryos (n=3 per condition), we processed the brain, stomach, liver and lung for H&E staining and immunofluorescence analysis to test the sensitivity of bioluminescence imaging to detect outgrowths. Within the 16 chick embryos injected with GFP-luc MCF10A RFP-TUBA1B cells, we observed no bioluminescence signal and, consistently, we found no tissues that contained GFP-positive cells (**Fig. 2e-g**)(**Supplementary Figure 5b**). In contrast, outgrowths were detected in 8 of the 33 chick embryos (24%) injected with GFP-AURKA/GFP-luc MCF10A RFP-TUBA1B cells (**Fig. 2e-g**); the sensitivity of outgrowth detection was similar for bioluminescence or immunofluorescence imaging. That is, we identified clusters of GFP-positive cells in luciferase-positive tissues taken from chick embryos injected with GFP-AURKA/GFP-luc cells (**Fig. 2h**). Taken together, our data shows that GFP-AURKA expression is sufficient to promote the survival, growth, and spread of non-cancer, human MCF10A RFP-TUBA1B breast cells when implanted in chicken embryos.

### Leader cells express AURKA during collective invasion

To study the effects of AURKA expression on cell migration, and assess the formation of leader cells during the process, we used in vitro scratch wound assays of MCF10A RFP-TUBA1B breast epithelial cells, which were imaged every 30 minutes. Leader cells were identified at 6 hours post-wounding by their position at the front of migrating groups and the presence of lamellipodium at their leading edge. Retrospective analysis of RFP-TUBA1B (microtubule) dynamics in leader cells (identified at 6h post-wounding) revealed random polarity of the centrosomes at wounding within future leader and their follower cells. Although random localization was maintained in follower cells through the time course, emerging leader cells polarized the centrosome to the front starting at 1-hour post-wounding, and were fully polarized by 3-hours, and this orientation was maintained as the wound closed (**Fig. 3a**). By assigning a numerical value to the centrosome position (front-polarized = 1.0; side-orientation = 0.5; and rear-orientation = 0), we observed significantly different kinetics of polarization in cells destined to be leader cells versus their follower cells (**Supplementary Fig. 6a**). Thus, active polarization of the centrosome distinguishes MCF10A breast epithelial cells destined to be leader cells from those destined to be follower cells.

**Figure 3.**
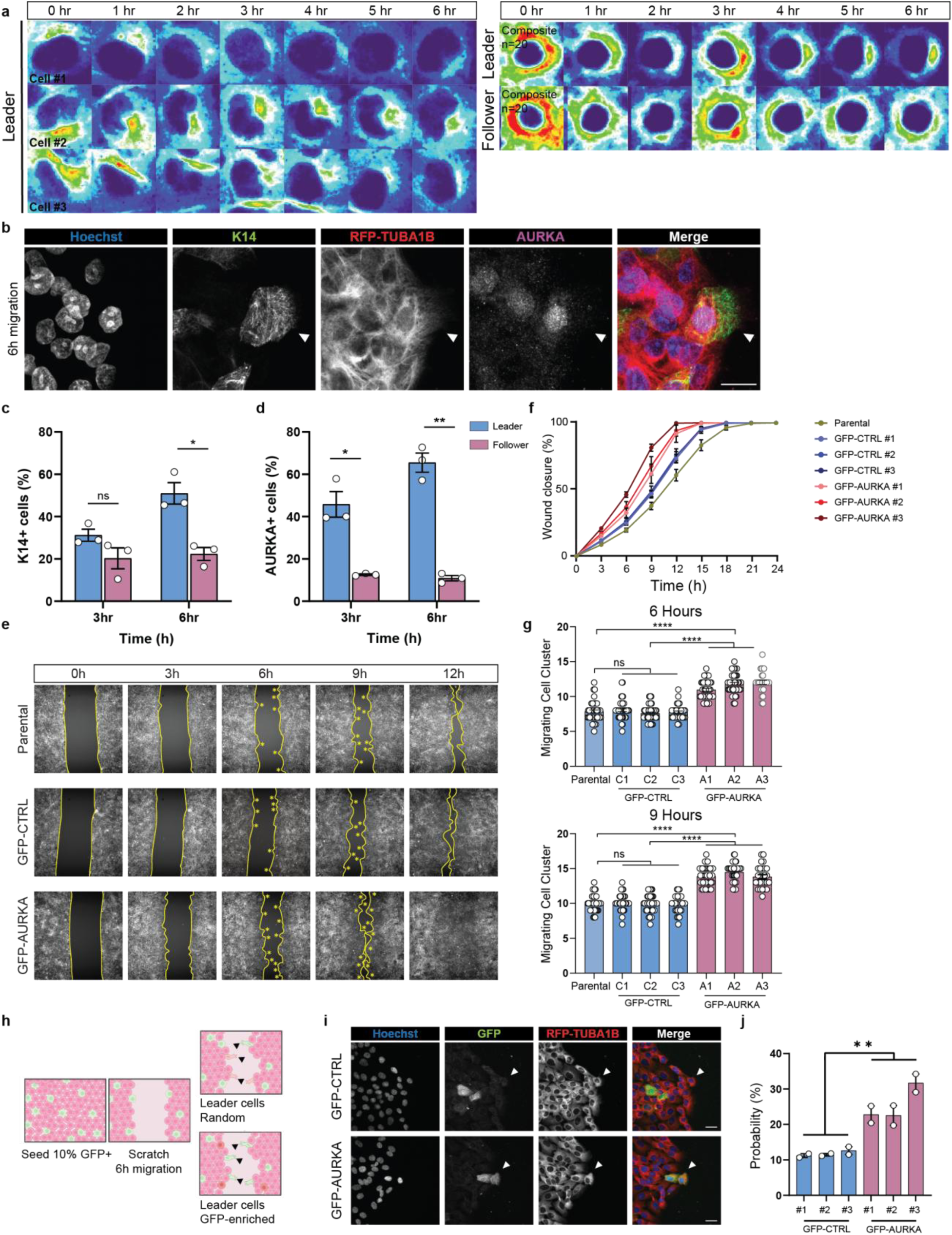
Leader cells express AURKA and GFP-AURKA promotes their emergence. **a.** Images from live-cell imaging of MCF10A RFP-TUBA1B scratch wound assay. Leader cells were identified at 6 hours by presence of lamellipodium and the dynamics of microtubule organization were retrospectively followed in leader cells and follower cells (left). Composite, overlay images of 20 leader or follower cells were generated by ImageJ (right). **b.** Representative images of a scratch wound assay of MCF10A RFP-TUBA1B cells. Cells were fixed six hours after wounding and stained with AURKA and K14. White arrowhead points to the leader cell. **c-d.** The proportion of leader and follower cells that are **(c)** K14-positive or **(d)** AURKA-positive at 3-hours or 6-hours post-wounding. Data are represented as mean ± SEM, n= 3 experiments. Student’s t-tests were used to determine significance. **e-g.** Scratch wound assay for parental (40 wounds), GFP-CTRL (#1=44, #2=42, #3=31 wounds), and GFP-AURKA (#1=43, #2=47, #3=27 wounds) cells imaged every 30 minutes over the course of 24 hours measuring **(f)** the percentage of wound closure or **(g)** the number of migrating cell groups (yellow stars) were quantified. Data are represented as **(f)** mean ± SEM or **(g)** mean ± SD, n= 3 experiments, P-value from multiple comparisons two-way ANOVA method t-test. **h.** Depiction of scratch wound assay with co-culturing of GFP-CTRL or GFP-AURKA with parental cells at a ratio of 1:10, growth for 48 hours until confluent, serum-starvation, and wounding. **i-j.** Cells were fixed at 6 hours post-scratch and stained with GFP. GFP-positive cells were frequently paired, indicating cell division that occurred post-seeding. Leader cells indicated by an arrowhead. **(j)** Probability of GFP-CTRL or GFP-AURKA cells becoming leader cells indicates a bias for GFP-AURKA cells. Data are represented as mean ± SEM, n= 2 experiments, P-value from Student’s t-test (n=6)

We postulated that centrosome polarization may correlate with expression of AURKA within cells destined to be leader cells. So, we examined the expression of a commonly used marker of leader cells, K14 [3], as well as AURKA, using IF analysis in cells fixed at 3 hours and 6 hours post-wounding (**Fig. 3b**). Leader cells were found to increase the expression of both K14 (**Fig. 3c**) and AURKA (**Fig. 3d**) following wounding (3h to 6h = AURKA-positive: 45.8% to 65.5%; K14-positive: 31.2% to 51%), whereas the proportion of AURKA-positive or K14-positive follower cells did not change with time (**Fig. 3c,d**). Similarly, the proportion of double-positive (AURKA^+^ K14^+^) leader cells almost doubled (3h to 6h = 13.1% to 25.0%) while double-negative (AURKA^-^K14^-^) leader cells decreased dramatically (3h to 6h = 36.1% to 8.5%) (**Supplementary Fig. 6b**). In follower cells, these proportions did not change, and most follower cells were double-negative (3h = 70.3%, 6h = 71.1%) (**Supplementary Fig. 6b**). Overall, these data suggest that AURKA expression correlates with a leader cell phenotype, including expression of K14.

### GFP-AURKA expression promotes the emergence of leader cells in vitro

To study the effects induced by elevated AURKA expression, we introduced GFP-control (GFP-CTRL) or GFP-AURKA into MCF10A RFP-TUBA1B parental cells (**Supplementary Fig. 2a,b**). We monitored scratch wound assays for parental, GFP-CTRL clones, and GFP-AURKA clones through live-cell imaging over the course of 24 hours (**Fig. 3e**). All three GFP-AURKA clones fully closed the wound faster than GFP-CTRL clones and parental cells (**Fig. 3f**). Moreover, the number of migrating cell groups per wound, which is indicative of the number of leader cells, were higher for all GFP-AURKA clones at both 6 hours and 9 hours post-wounding (**Fig. 3g**). Next, we investigated whether GFP-AURKA cells have a higher propensity to become leader cells during wound closure. To do so, we used a competitive co-culture approach. GFP-CTRL or GFP-AURKA cells were seeded at a ratio of 1:10 with parental MCF10A RFP-TUBA1B cells (**Fig. 3h**). After seeding, cells were cultured for 24 hours to become confluent, and then serum-starved prior to scratch wound assays. Wounds were fixed at 6 hours post-wounding and GFP-positive leader cells were identified (**Fig. 3i**). These competition assays indicated an enrichment for all three GFP-AURKA clones to become leader cells compared to GFP-CTRL clones (**Fig. 3j**). Thus, a majority of leader cells are AURKA-positive, and expression of GFP-AURKA increases the number of leader cells in a wound, and is sufficient to endow transfected cells with a heightened propensity to become a leader cell.

### AURKA activity suppresses cell scattering and is needed for collective invasion

To determine the necessity of AURKA activity for leader cell formation, we suppressed AURKA activity using the small-molecular inhibitor MLN8237 (aka Alisertib). First, we performed a dose-response assay exposing MCF10A cells to a variety of MLN8237 doses and, through immunofluorescence detection of phosphorylated AURKA, determined inhibitor efficacy (**Supplementary Fig. 7a**). Then, we treated scratch wound assays using MCF10A RFP-TUBA1B cells, and found that addition of increasing concentrations of MLN8237 resulted in a decrease in wound closure and a decrease in the number of migrating cell groups per wound in a dose-dependent manner over the course of 24 hours (**Fig. 4a-c**). Interestingly, cells scattered, or separated from the main monolayer of cells, and this phenotype increased in a dose-dependent manner after the addition of MLN8237 (**Fig. 4d**) suggesting AURKA activity might be crucial for the cohesiveness of migrating cell groups. So, we tracked the migration pattern of leader cells in the presence of MLN8237, or DMSO, and found that MLN8237-treated leader cells displayed both a loss of directionality (**Fig. 4e**) and a loss of velocity (**Fig. 4f**).

**Figure 4.**
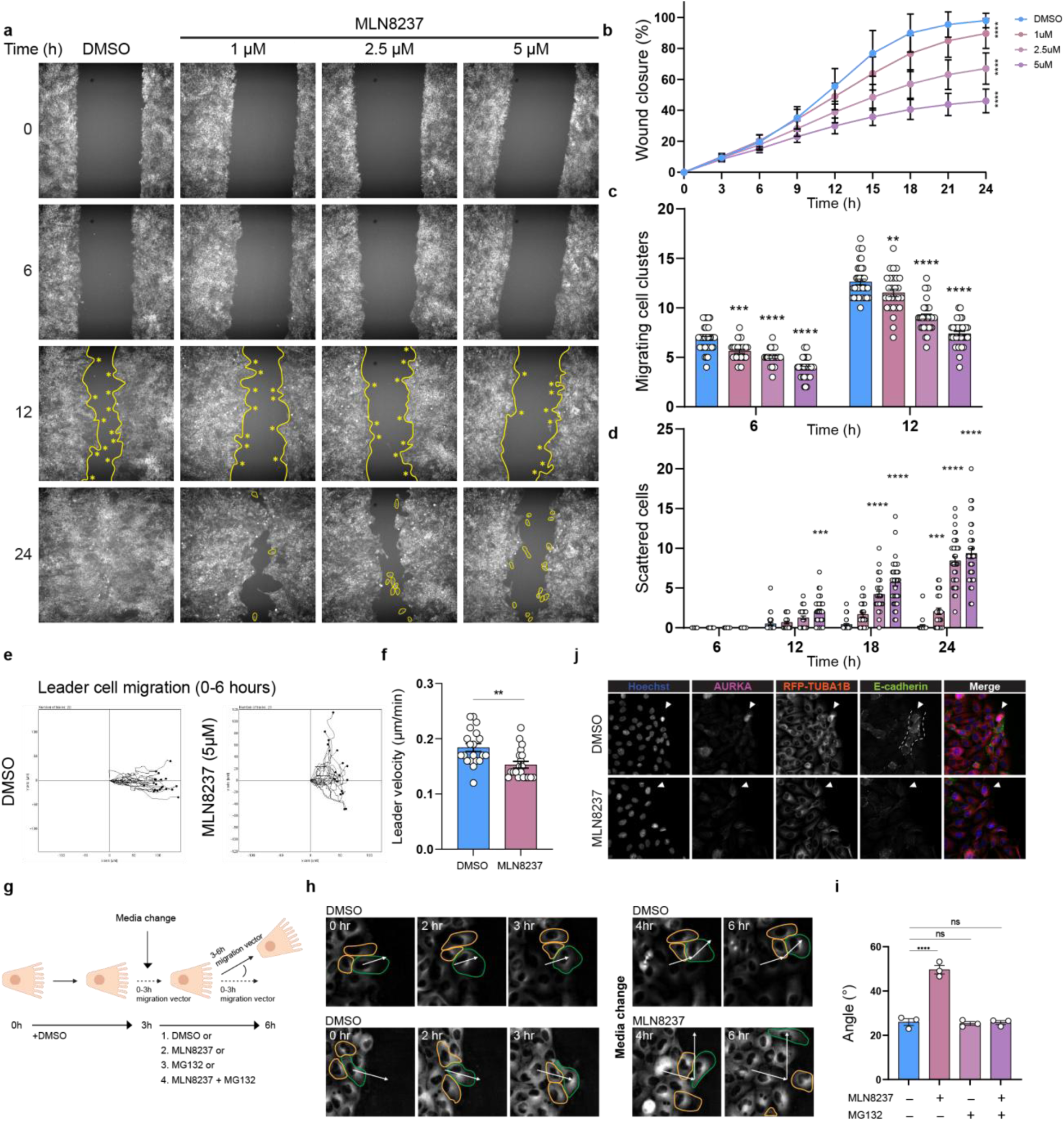
AURKA activity suppresses cell scattering and is needed for collective cell migration. **a.** Live-cell imaging of scratch wound assays treated with DMSO or MLN8237 (1μM, 2.5μM, 5μM). Images were taken every 15 minutes over the course of 24 hours. At 12 hours, the wound is outlined by a yellow line and migrating cell clusters are marked by an asterisk (*). Scattered cells are highlighted with a yellow contour. **b-d.** Live-cell images were used to quantify **(b)** the wound closure speed **(c)** migrating cell clusters and **(d)** scattered cells for cells treated with DMSO (36 wounds) or MLN8237 at a concentration of 1μM (27 wounds), 2.5μM (35 wounds) and 5μM (36 wounds). Data are represented as mean ± SD, n = 3 independent experiments, P-value from multiple comparisons two-way ANOVA method t-test. **e.** The migration of leader cells with or without the addition of MLN8237 (5 μM) at 0 hours post-scratch were tracked. Leader cells were identified retrospectively at 6 hours using 10x objective images. **f.** Velocity of leader cells with or without the addition of MLN8237 (5 μM) were determined. **g.** Schematic of scratch wound assay with or without treatment of MLN8237 or MG132. After making the scratch, media with DMSO was added and cells were allowed to migrate. At 3 hours post-scratch, media with or without treatment of MLN8237 (5μM) or MG132 (1 μM) was added. **h.** Live-cell imaging was used to observe scratch wound assays treated with or without MLN8237 (5 μM) for 6 hours post-scratch. Leader cells are outlined in green, and two follower cells are outlined in orange. White arrows represent migration vector of the leader cell. **i.** Quantification of the difference between the migration vector from 0-3 hours and 3-6 hours in response to MLN8237 or MG132. Data are represented as mean ± SEM, n = 3 independent experiments, P-value from multiple comparisons two-way ANOVA method t-test. **j.** IF analysis of AURKA and E-cadherin expression in migrating MCF10A RFP-TUBA1B cells. The migrating cell cluster have a leader cell (white arrowhead) with high AURKA expression and followers that are positive for E-cadherin. Intercellular bridges between cells at the edge of the wound are also E-cadherin positive.

Scattering occurred within hours of exposure to MLN8237, suggesting rapid disruption of cell cohesion. So, we hypothesized this rapid scattering was proteasome-dependent and, consequently, may be reversible in the presence of a proteasome inhibitor, MG132. To test this, we performed a scratch in the presence of DMSO and imaged wound closure for 3 hours, to allow for the formation of leader cells with directional migration. At 3 hours, the media was changed with the addition of DMSO, MLN8237, MG132, or a combination of both, and wound closure was imaged for a further 3 hours (**Fig. 4g**). To measure the direction of migration, and cell scattering from that direction, we determined a vector of migration for leader cells identified at 3-hours post-wounding as well as a vector of migration for those same leader cells between 3- and 6-hours post-wounding. In DMSO-treated cultures, the variance between the two vectors was small (<30^⸰^) (**Fig. 4h,i**). However, treatment with MLN8237 (5μM) resulted in significant deviance from the expected vector of migration, and follower cells quickly detached from the group post-treatment. Inclusion of MG132, however, was sufficient to rescue both effects of MLN8237, and normalize the vector of leader cell migration and the cohesion of follower cells (**Fig. 4i**).

To assess the needed role for AURKA in the maintenance of cell cohesion, we first examined E-cadherin-positive contacts in follower cells. In wounds fixed at 6-hours post-wounding, we noted that E-cadherin-positive islands of follower cells trailed AURKA-positive leader cells (**Fig. 4j**) often including very thin E-cadherin-positive intercellular bridges connecting the leader cell with neighbouring cells. Addition of MLN8237 (5 μM for 3 hours) decreased the E-cadherin intensity between the leader cell and follower cells at the wound edge (**Fig. 4j**) (**Supplementary Figure 7b,c**).

### AURKA locates EPLIN to E-cadherin-positive cell-cell contacts in follower cells

Migrating leader cells transport lysosomes along microtubules to accumulate at their leading edge, which promotes leader cell formation and migration [28]. Because MG132 treatment can alter lysosomal activity [29, 30], we next followed lysosome positioning in living wounds. MCF10A RFP-TUBA1B cells were wounded and allowed to invade for 4 hours before incubating cells with tubulin and lysosome dyes for 2 hours. Lysosomes localized to microtubules and were most concentrated at the centrosome microtubule bundle (**Supplementary Fig. 8a**). We added DMSO, MLN8237, MG132 or both MLN8237 and MG132 (Dual) and quantified the ratio of lysosomes localized around the centrosome region compared to the whole cell (**Supplementary Fig. 8b**). MLN8237 and MG132 separately led to a non-significant increase in lysosome scattering after 2 hours, concurrent with leader cell scattering; however, dual treatment did not rescue lysosome scattering (**Supplementary Fig. 8c,d**), indicating that the disruption of lysosome positioning along microtubules may not explain how AURKA inhibition induces leader cell scattering in a MG132-dependent manner.

Next, we examined a published AURKA interactome dataset [31] and a published AURKA phosphoproteome dataset [32] and identified 17 overlapping gene products that have both the potential to interact with and be a substrate for AURKA. Only one gene/gene product, termed *LIM domain and Actin binding 1* or *LIMA1*, was found to localize to adherens junctions (**Fig. 5a**). *LIMA1* encodes epithelial protein lost in neoplasia (EPLIN), an actin-binding protein that stabilizes the cadherin-catenin complex at cell contacts [33]. And, we confirmed the physical complex formation between AURKA and EPLIN through co-precipitation of EPLIN in AURKA immunoprecipitates (**Fig. 5b**)(**Supplementary Fig. 9**). Next, we performed immunofluorescence analysis of EPLIN and E-cadherin in migrating MCF10A cells and found that EPLIN and E-cadherin both localize at the intercellular bridges (**Fig. 5c**). MLN8237 treatment, however, reduced EPLIN intensity at intercellular bridges (**Fig. 5d**). In fact, we found a dramatic localization of EPLIN to the lamellipodia at the leading edge upon the inhibition of AURKA (**Fig. 5e**). So, we next followed cytoskeleton dynamics, including microtubules (SPY555-tubulin and RFP-TUBA1B) and actin (SPY650-FastAct), during scratch wound assays performed in the presence (+DMSO) (**Supplementary Fig. 10**) or absence (+MLN8237) (**Supplementary Fig. 11**) of AURKA activity. To do so, MCF10A RFP-TUBA1B cells were wounded and allowed to invade the wound for 2 hours before incubating cells with SPY555-tubulin and SPY650-FastAct for 1 hour. Then, wounds were incubated with MLN8237 or DMSO control followed by live imaging at 10- or 20-minute intervals. In DMSO-treated cultures, front-polarized centrosomes nucleated microtubule networks directed to the leading edge, which overlapped with actin networks at lamellipodia (**Fig. 5f**); these co-localizing networks correlated with directional cell migration into the wound (**Fig. 5f**). However, MLN8237 treatment led to the realignment of the microtubule cytoskeleton, including loss of centrosome nucleation, coincident with an alteration in the actin network and a redirection of the leader cell (**Fig. 5f**) detaching from their followers (**Supplementary Figs. 10,11**). Thus, we conclude that AURKA activity is required for directional migration in leader cells through the concurrent polarization of the microtubule network at the centrosome and the actin network at lamellipodia, which correlates with the subcellular location of EPLIN, an actin-binding protein that co-localizes with E-cadherin+ cell-cell contacts conditional upon AURKA activity.

**Figure 5.**
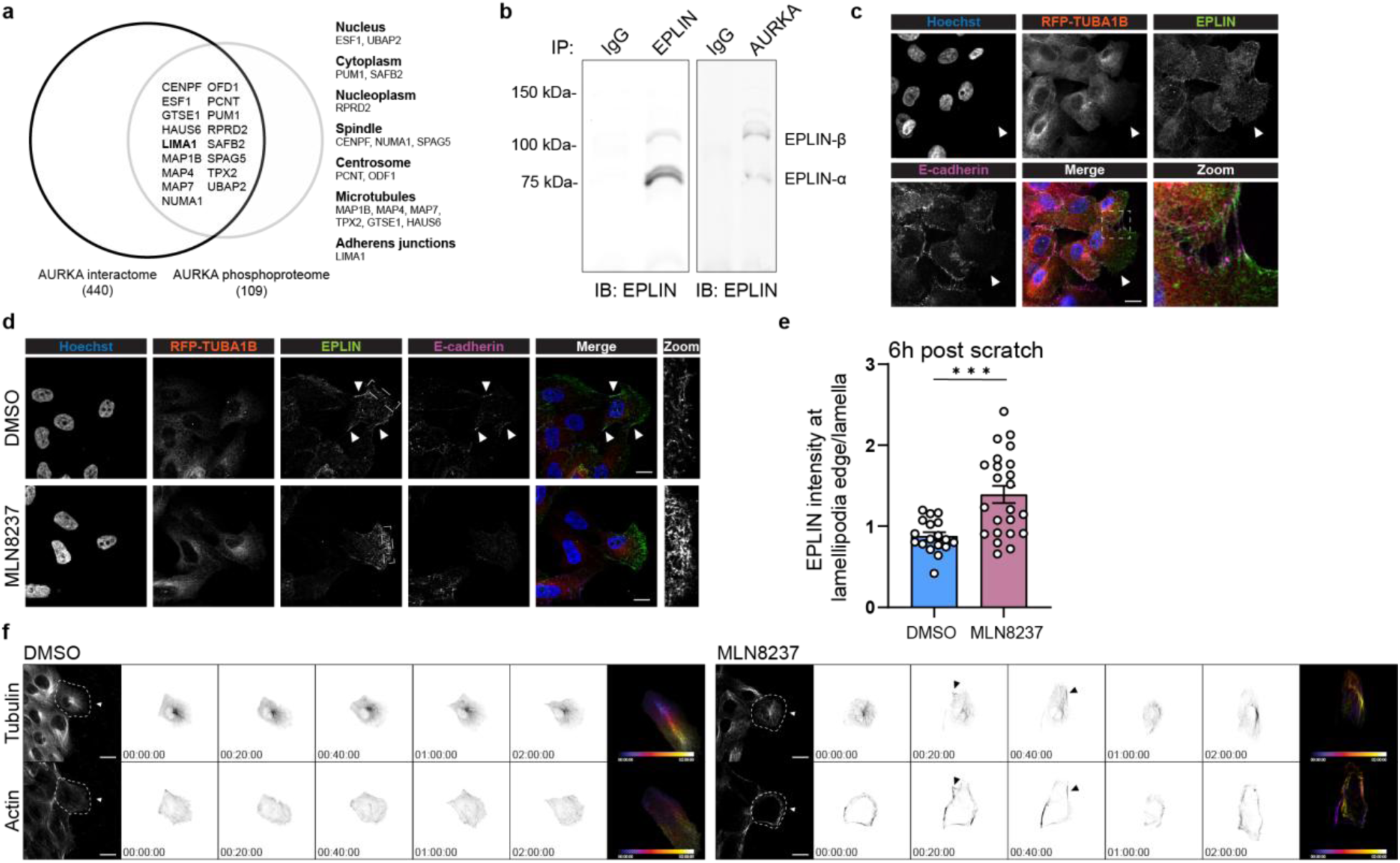
AURKA locates EPLIN to E-cadherin-positive cell contacts to maintain follower cell cohesion. **a.** Venn diagram showing 17 overlapping genes between published datasets of the AURKA interactome [31] and the AURKA phosphoproteome [32]. **b.** Immunoprecipitation of MCF10A migration-enriched cells using EPLIN (left) or AURKA (right) antibodies, then blotted with EPLIN antibody. **c.** Super-resolution microscopy images of leader cell with follower cells connected with EPLIN and E-cadherin positive intercellular bridges (zoom image). **d.** Super-resolution microscopy images EPLIN and E-cadherin cell contacts and relocalization of EPLIN to lamellipodia following exposure to MLN8237. **e.** DMSO (18 cells) or MLN8237 (23 cells) treated scratch wounds were analyzed for EPLIN intensity at the lamellipodia edge/lamella at 6 hours post-scratch. P values were calculated by Student’s t-test. **f.** Live-cell imaging of scratch wound treated with DMSO or MLN8237 using SPY555-tubulin and SPY650-FastAct dye to stain microtubules and actin. White arrowheads point to leader cells. Black arrowheads point to altered and aligned microtubule and actin cytoskeleton in MLN8237-treated cells. Dotted region represents leader cell mask used to center inverted images. Temporal color-coded image of microtubules and actin within a leader cell is shown for 2-hour imaging.

## Discussion

The metastatic process is believed to initiate with collective cell invasion into surrounding tissues. Our examination of public datasets reveals enrichment of “G2/M” and “Mitotic spindle” signatures (strongly correlated with *AURKA* expression) at metastatic sites, and strong positive correlation between *AURKA* expression and metastatic signatures in pan-cancer datasets. Collective invasion is often guided by leader cells, which are linked via E-cadherin-positive contacts and actomyosin cables to the migratory group of followers [5, 6]. Here, we discovered that GFP-AURKA expression in a non-malignant human breast cell-line is sufficient to promote engraftment and endow outgrowths within the chicken embryo CAM model. Live imaging of scratch wound assays established GFP-AURKA expression to be sufficient, and AURKA activity to be necessary, to endow a leader cell phenotype; in the absence of AURKA activity, individual cells disperse in migrating wounds due to the dissolution of cell adhesion with follower cells, and resulting in a dramatic re-organization of the actin cytoskeleton and its regulator Epithelial protein lost in metastasis (EPLIN).

In migratory leader cells, front-rear polarity involves front positioning of the centrosome, a microtubule organizing centre, and the protrusion of actin-rich lamellipodia at the leading edge; consequently, interconnection of actin and microtubule cytoskeletal networks is critical to maintaining this phenotype. Mechanisms of interconnection are diverse, including mechanical and biochemical signals that influence RhoA activity via direct mechanical traction force [34], or through pushing force from follower cells [35], and through biochemical communication often delivered by the microtubule cytoskeleton, such as the motor kinesin-1 modulating the activity of RhoA, via the availability of a RhoA GEF [36], or the directional delivery and accumulation of lysosomes at the leading edge to regulate local Rac1 activity [28]. Here, we add to these mechanisms with the demonstration that AURKA activity is essential, and its inhibition has immediate consequence on the positioning of the centrosome and the organization of microtubule and actin cytoskeletal networks, which results in the dissolution of cell contacts and the scattering of follower cells.

AURKA regulates the microtubule cytoskeleton during cell division but it has additional functions in non-mitotic cells [20]. For example, AURKA has non-kinase activity in the nucleus, which transactivates the *MYC* promoter to enhance stem-like phenotypes [37, 38]. AURKA is also needed for an EMT phenotype in carcinoma cells [39, 40], including during the collective invasion of breast carcinoma cells [41]. Mechanistically, AURKA regulates the microtubule cytoskeleton in quiescent cells, such as through the disassembly of the primary cilium, via NEDD9-HDAC6 [42], and the promotion of neurite extension in neurons [43]. Indeed, NEDD9 binds to cortactin, leading to its deacetylation in an AURKA/HDAC6 manner, with direct impact on actin dynamics in lamellipodium and their protrusion [44]. AURKA phosphorylates S296 NEDD9 to promote its degradation [45], and elevated expression of NEDD9 disrupts E-cadherin through lysosomal degradation [46–48]. Consistently, we find that E-cadherin junctions are immediately disrupted following inhibition of AURKA activity, which induces a scattering phenotype.

Given the recent evidence that lysosome accumulation along microtubules regulate leader cell emergence through the promotion of lamellipodia [28], we performed live cell imaging experiments to measure lysosome abundance at centrosome regions in leader cells. We confirmed that lysosomes accumulate along microtubules and find that dual AURKA and proteosome inhibition reduces lysosome accumulation within these centrosome regions. Furthermore, we observed a dramatic localization of EPLIN to the lamellipodia following inhibition of AURKA activity. EPLIN is a multifunctional protein that is critical to the connection of the actin cytoskeleton and adherens junctions [49], and its localization to lamellipodia mechanistically explains the dissolution of these junctions, possibly in an additive manner with lysosomal degradation. Additional studies are required, however, to identify the specific EPLIN sites that may be phosphorylated by AURKA. Moreover, future research is required to uncover additional, putative molecular mechanisms because AURKA-mediated control of microtubule and centrosome dynamics is likely to have multifactorial consequences.

Our results indicate that AURKA may be a particularly robust candidate for targeting during metastatic dissemination; along with *ESR1* and *ERBB2* expression, *AURKA* expression is sufficient to categorize breast cancer subtypes [13], and we find its expression is highly correlated with pan-cancer metastatic dissemination. Moreover, the results of a recent phase 2 study of Alisertib (MLN8237), with and without fulvestrant, indicate promising clinical activity for Alisertib monotherapy in patients with CDK4/6 inhibitor-resistant or endocrine-resistant metastatic disease [50]. However, it will be essential to understand and balance the inhibitor’s activity against tumor cells with the potential for off-target effects on non-cancer cells, such as reduced anti-tumor immunity [51, 52]. In conclusion, our data demonstrate that *AURKA* expression is sufficient to enable collective invasion and metastasis making its inhibition a potential therapeutic strategy to prevent metastasis of human breast carcinoma cells.

## Acknowledgements

This study was partially funded by the Canadian Institutes of Health Research (CIHR F-19-03865, TFRI PPG F22-00533, CIHR F22-03789, CIHR F24-00975) and a Canadian Cancer Society Challenge grant (grant #709031). BPZ received a Michael Cuccione Foundation for Childhood Cancer Research studentship. TLHC received a Canadian Breast Cancer Foundation graduate studentship. AG Received a CIHR Canadian Graduate Scholarship – Masters. RG received a Canadian Cancer Society Research Training Award – PhD.

## Author contributions

These authors contributed equally: Bo Peng Zhou, Tony LH Chu, Aidan Gallant, and Ryan Ghorayeb

BPZ: investigation, data curation, formal analysis, writing - revised draft. TLHC: investigation, data curation, formal analysis, writing – original draft. AG: investigation, data curation, writing – revised draft. RG: investigation, data curation, formal analysis, writing – revised submission, SW: investigation – chicken CAM model. TB: investigation – chicken CAM model. CG: investigation, data curation. REG: formal analysis. CJL: supervision, investigation – chicken CAM model, project administration. MAP: supervision, formal analysis. CAM: conceptualization, funding acquisition, supervision, project administration, writing draft.

## Data availability statement

All other relevant data supporting the key findings of this study are available within the article and its Supplementary Information files or from the corresponding author upon request. Source data are provided with this paper.

