## supplemental figures for "Aurora kinase A enables collective invasion and metastasis by endowing a leader cell phenotype and stabilizing cohesion with follower cells"

Supplementary Information

### Supplementary Figures

a

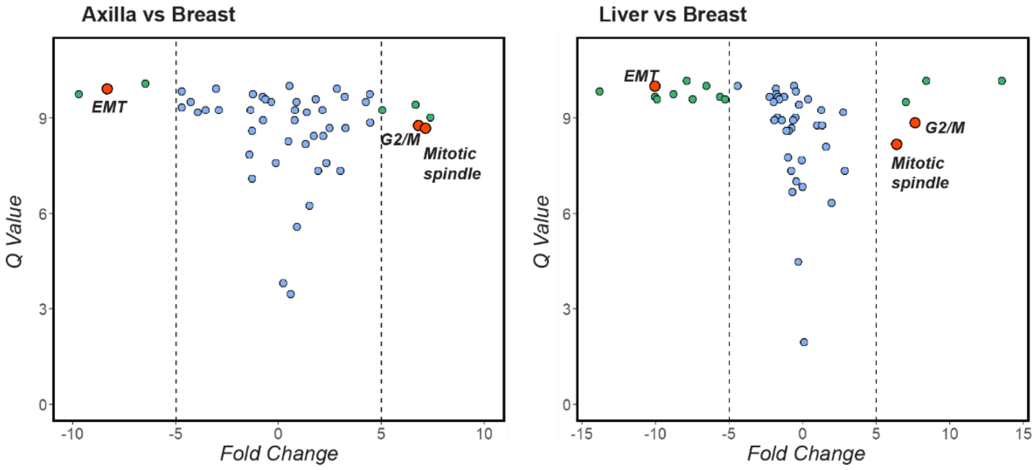


b

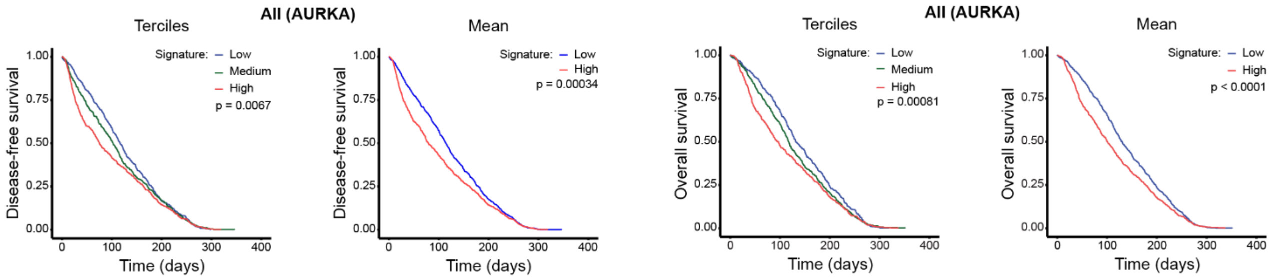


c

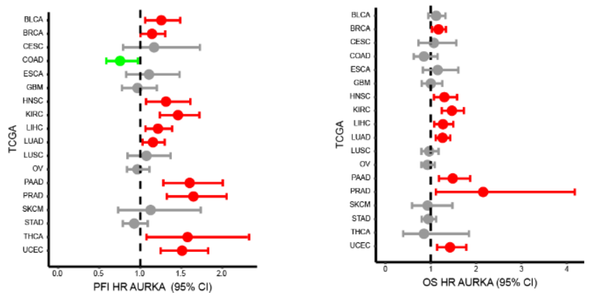


**Supplementary Figure 1. High *AURKA* expression is associated with worse prognosis in breast cancer patients in METABRIC cohort.**

**a.** Scatter plot of Q value versus Fold Change for Axilla versus Breast and Liver versus Breast for each Hallmark Gene Set. Gene sets (green) and pathways of interest (red) that are both significantly different and have an absolute fold change value of 5 or more are highlighted. Data sourced from [9].

**b.** Kaplan-Meier plots with rank test p-values are shown for the association between *AURKA* expression and disease-free survival (DFS) and overall survival (OS) for all breast cancer subtypes in terciles (left) or mean (right), respectively.

**c.** Forest plot displaying the hazard ratio (HR) for overall survival (OS) or progression-free interval (PFI) of AURKA expression across TCGA studies.

**a**

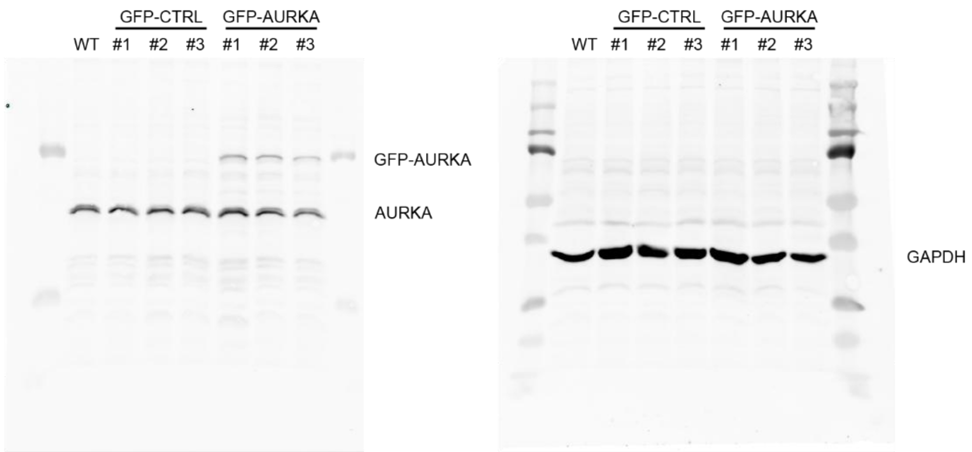


**b**

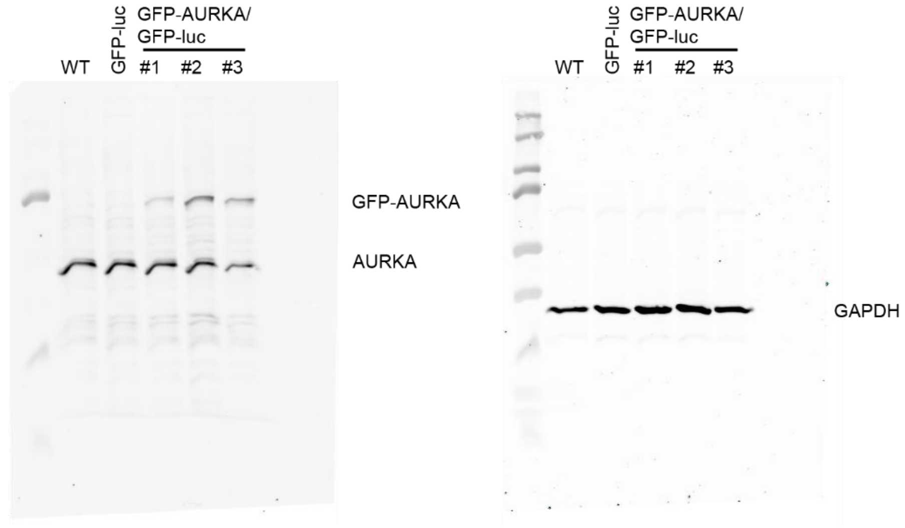


**Supplementary Figure 2. Western blot analysis of MCF10A RFP-TUBA1B clones.**
**a.** Western blot analysis of parental MCF10A RFP-TUBA1B (WT), and clones expressing GFP-CTRL or GFP-AURKA, respectively. GAPDH served as a loading control.
**b.** Western blot analysis of clones expressing GFP-luc or GFP-AURKA/GFP-luc, respectively. GAPDH served as a loading control.

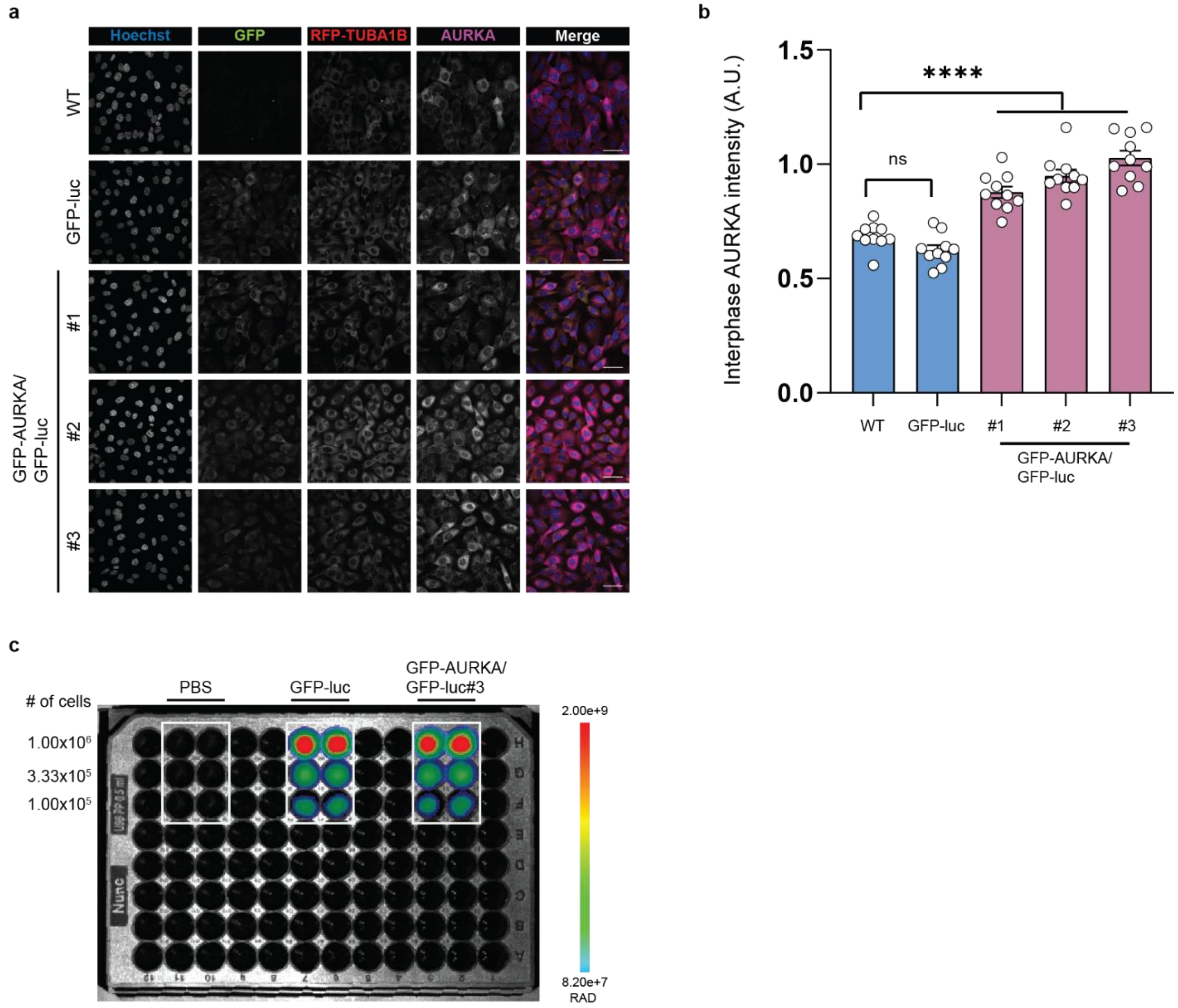


**Supplementary Figure 3. Characterization of GFP-luciferase expression in MCF10A RFP-TUBA1B cells.**

**a.** Immunofluorescence analysis of interphase parental MCF10A RFP-TUBA1B (WT), and cells expressing GFP-luc or GFP-AURKA/GFP-luc, respectively. Imaging was performed on a confocal microscope.

**b.** The intensity of AURKA immunofluorescence was measured in arbitrary units (AU) within interphase cells. n=2 experiments (5 random regions for each cell per experiment).

**c.** Bioluminescence imaging of GFP-luc or GFP-AURKA/GFP-luc #3 clones, seeded at specified cell number titrations, was measured using IVIS, with PBS-loaded wells serving as a negative control.

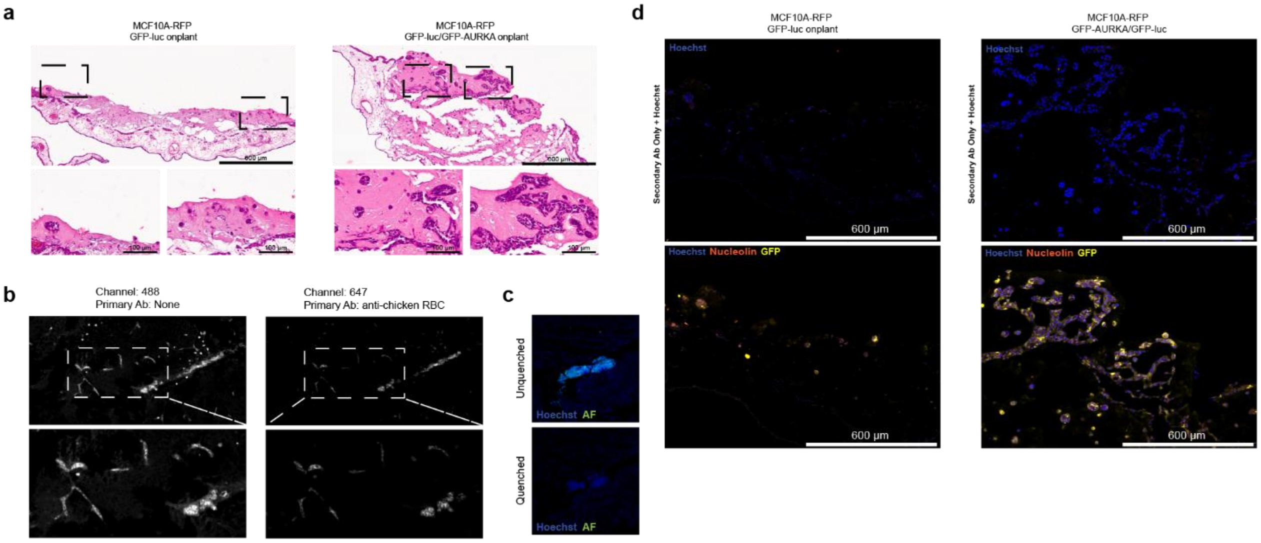


**Supplementary Figure 4. GFP-AURKA expression enhances engraftment in a chicken embryo model.**

**a.** H&E staining of formalin-fixed, paraffin-embedded CAM tissues isolated at day 6 following engraftment.

**b.** Chicken red blood cells, both young and adult, have a nucleus and have autofluorescence in the Alexa 488 channel.

**c.** To reduce non-specific autofluorescence of red blood cells, tissues were quenched using a commercial kit.

**d.** Immunofluorescence detection of GFP and a human-specific nucleolin was used to confirm the presence of human cells. Because of potential autofluorescence in the Alexa 488 channel, secondary antibody only was included as a negative control.

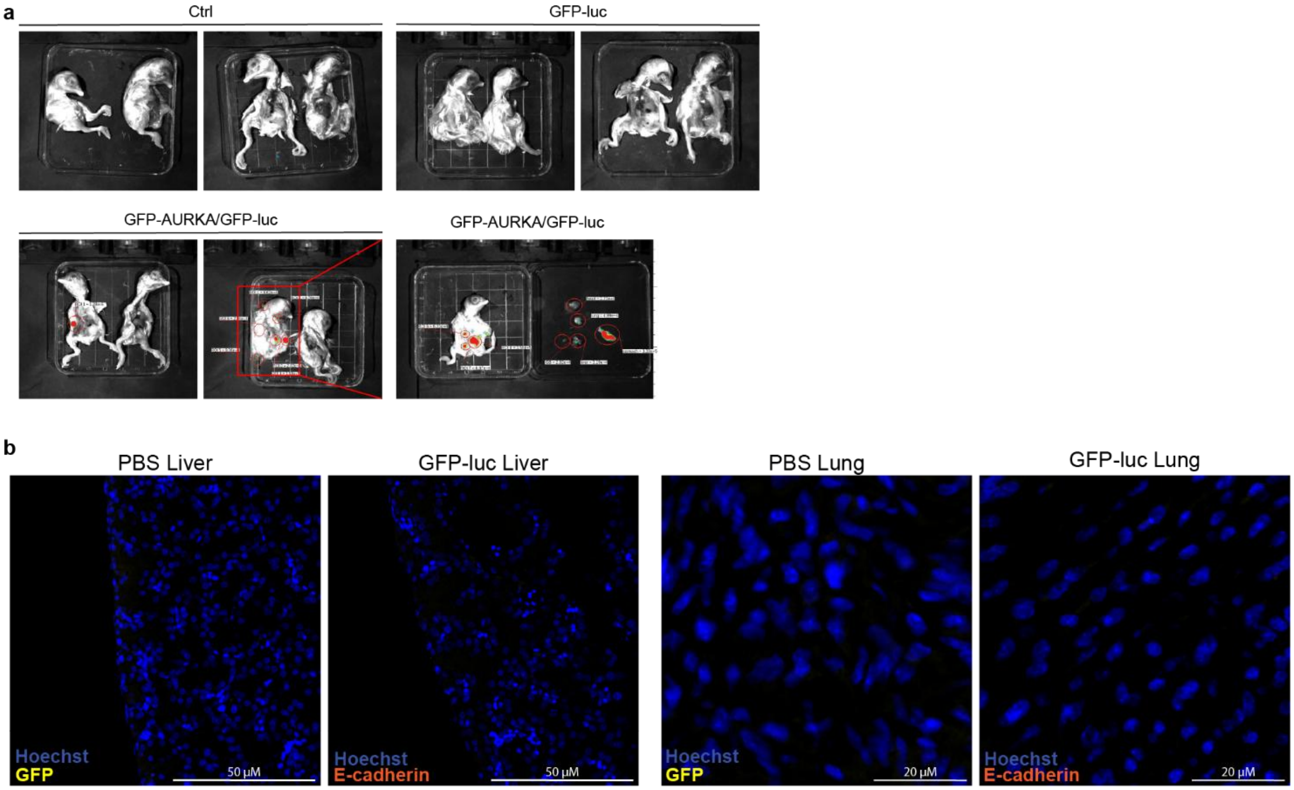


**Supplementary Figure 5. GFP-AURKA expression enhances metastatic potential in a chicken embryo model.**

**a.** Representative IVIS images of control (PBS injected), GFP-luc injected, and GFP-AURKA/GFP-luc injected chick embryos.

**b.** Immunofluorescence analysis of CAM tissues, previously injected with PBS or GFP-luc cells, stained with GFP or E-cadherin.

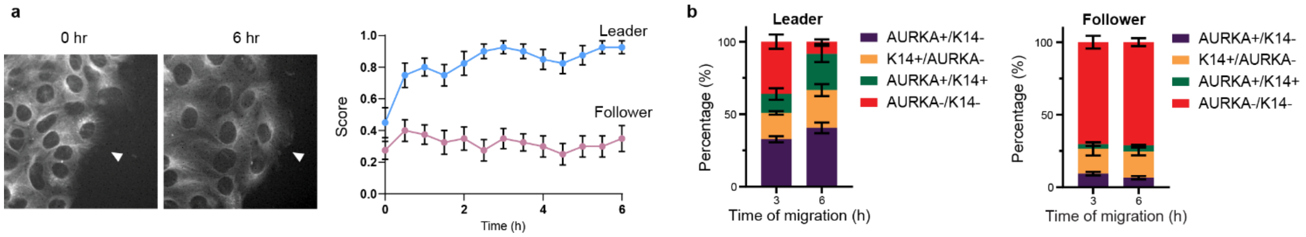


**Supplementary Figure 6. Leader cells front-polarize their centrosome and express AURKA.**

**a.** Scoring scheme to monitor centrosome polarity of leader cells (green) and follower cells (red). Front polarization: 1 point, side polarization: 0.5-point, rear polarization: 0 point. Centrosome polarization score is plotted for leader cells and follower cells over 6 hours scratch wound assay, n = 20 cells.

**b.** The proportion of leader and follower MCF10A RFP-TUBA1B cells that are AURKA+ only, K14+ only, double positive, or double negative. Data are represented as mean ± SEM, n= 3 experiments.

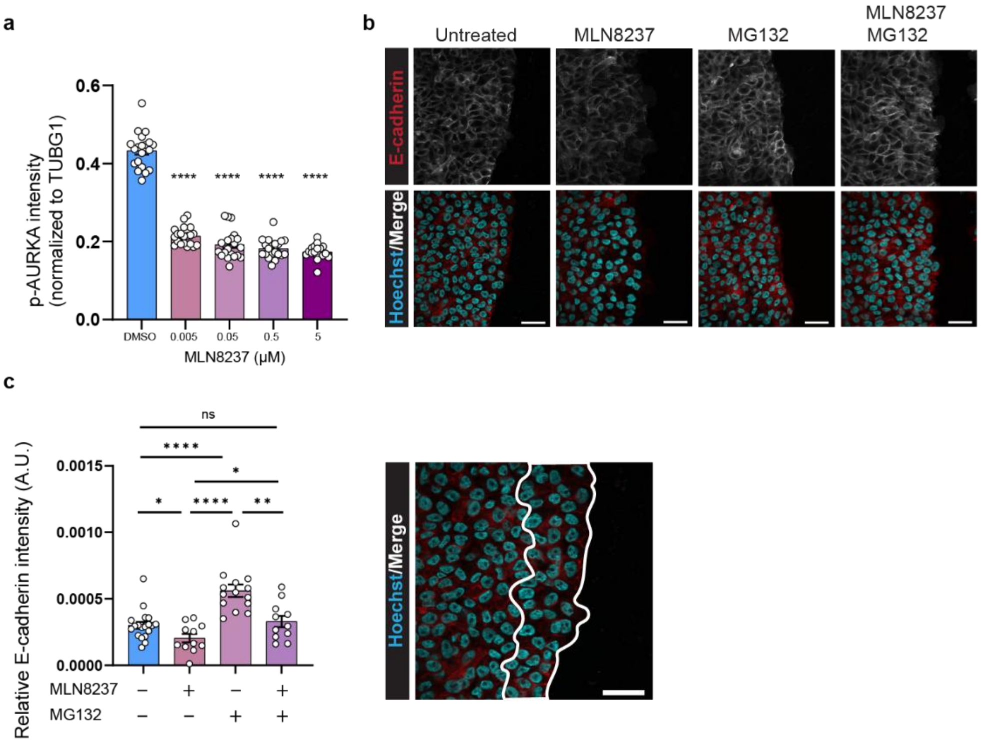


**Supplementary Figure 7. AURKA activity maintains follower cell cohesion.**
**a.** IF analysis of phosphorylated AURKA (p-AURKA) shows a reduced intensity with increasing concentration of MLN8237.
**b.** Scratch wound assay treated with or without MLN8237 (5uM) or MG132 (1 uM) (DMSO = 19, MLN8237 = 11, MG132 = 13, MLN8237 + MG132 = 11, from two independent experiments) fixed at 6 hours and stained for E-cadherin.
**c.** E-cadherin intensity of migrating cells were quantified by outlining the wound edge cells (outline in white).

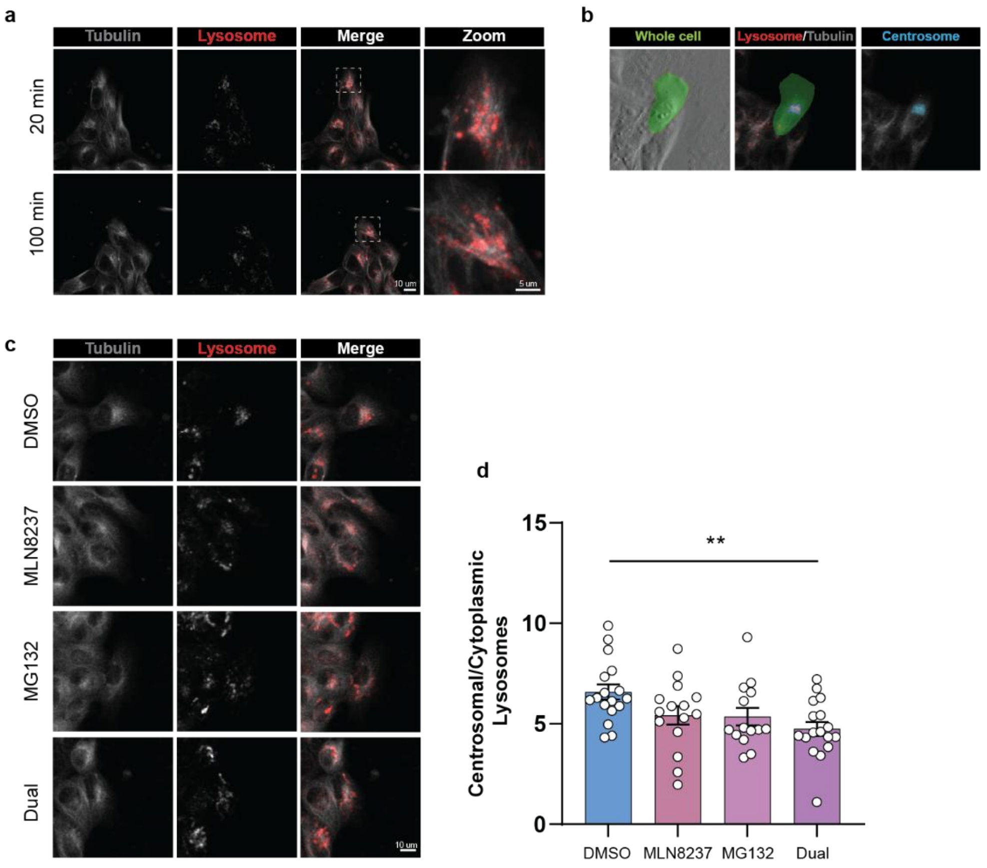


**Supplementary Figure 8. Dual AURKA and proteasome inhibition alters lysosome abundance at centrosome regions in leader cells.**

**a.** Live-cell imaging of MCF10A RFP-TUBA1B cells using both SPY555-tubulin and LysoTracker™ dyes.

**b.** Live-cell images of MCF10A RFP-TUBA1B cells using both SPY555-tubulin and LysoTracker™ dyes were divided into two regions, whole cell drawn by the phase contrast channel and centrosome region which was defined by 3-sigma thresholding bright spot algorithm for the tubulin channel.

**c.** Live cell imaging of MCF10A RFP-TUBA1B cells treated with DMSO, MLN8237, MG132, or dual treatment.

**d.** Quantification of lysosome abundance in centrosome regions compared to whole cell. Data points are individual cells from n=4 experiments, significance was determined via one-way ANOVA with Tukey’s post hoc test.

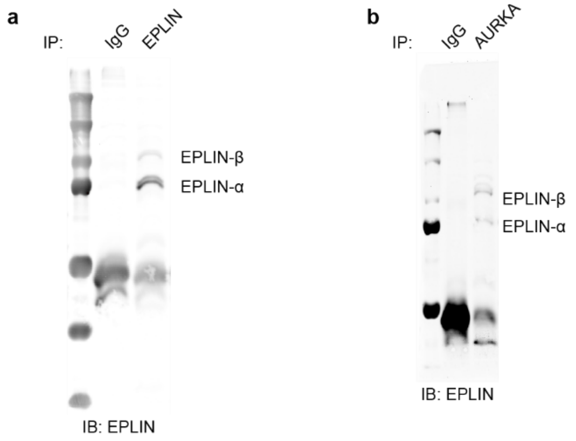


**Supplementary Figure 9. Co-immunoprecipitation of AURKA and EPLIN.**
Cell lysates were collected by growing cells in a 10 cm dish, then scratched in a grid pattern to enhance the number of migrating cells. Cells were collected 12 hours post-scratch. Cell extracts were used for immunoprecipitation using Dynabeads and an **(a)** anti-EPLIN or IgG control, or and **(b)** anti-AURKA antibody or IgG control. Resulting IP sample was used to run SDS-PAGE and stained for EPLIN.

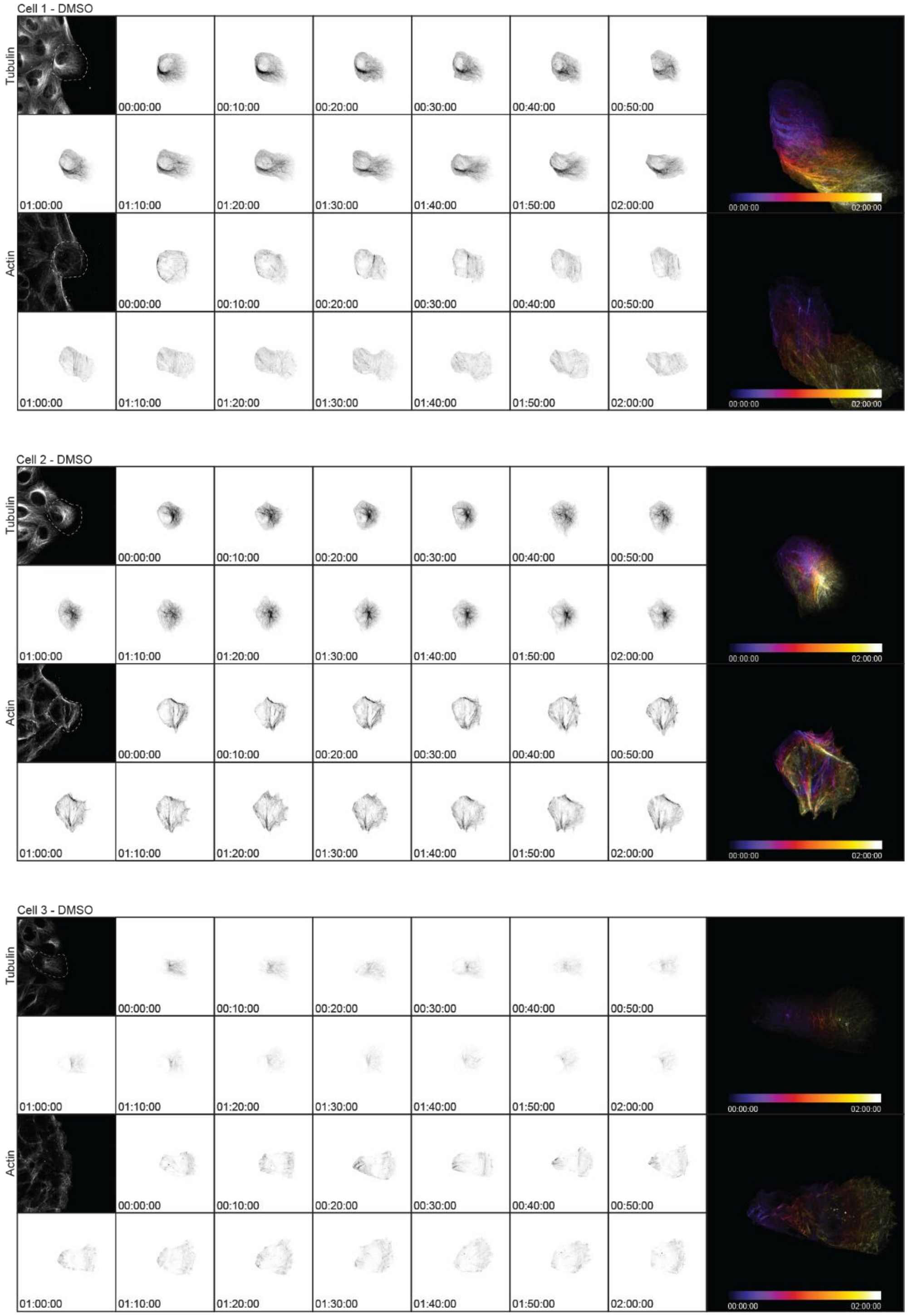


**Supplementary Figure 10. Live-cell imaging of tubulin and actin in MCF10A RFP-TUBA1B leader cells treated with DMSO.** Live-cell imaging of MCF10A RFP-TUBA1B cells using SPY555-tubulin and SPY650-FastAct. After the addition of DMSO, cells were monitored for 2 hours at a time interval of 10 minutes. Inverse images of cropped leader cells, then centered around the nucleus (crop is highlighted in dashed lines in the raw images). Time-color coded overlay images of non-centered, cropped leader cells are shown for 2 hours of

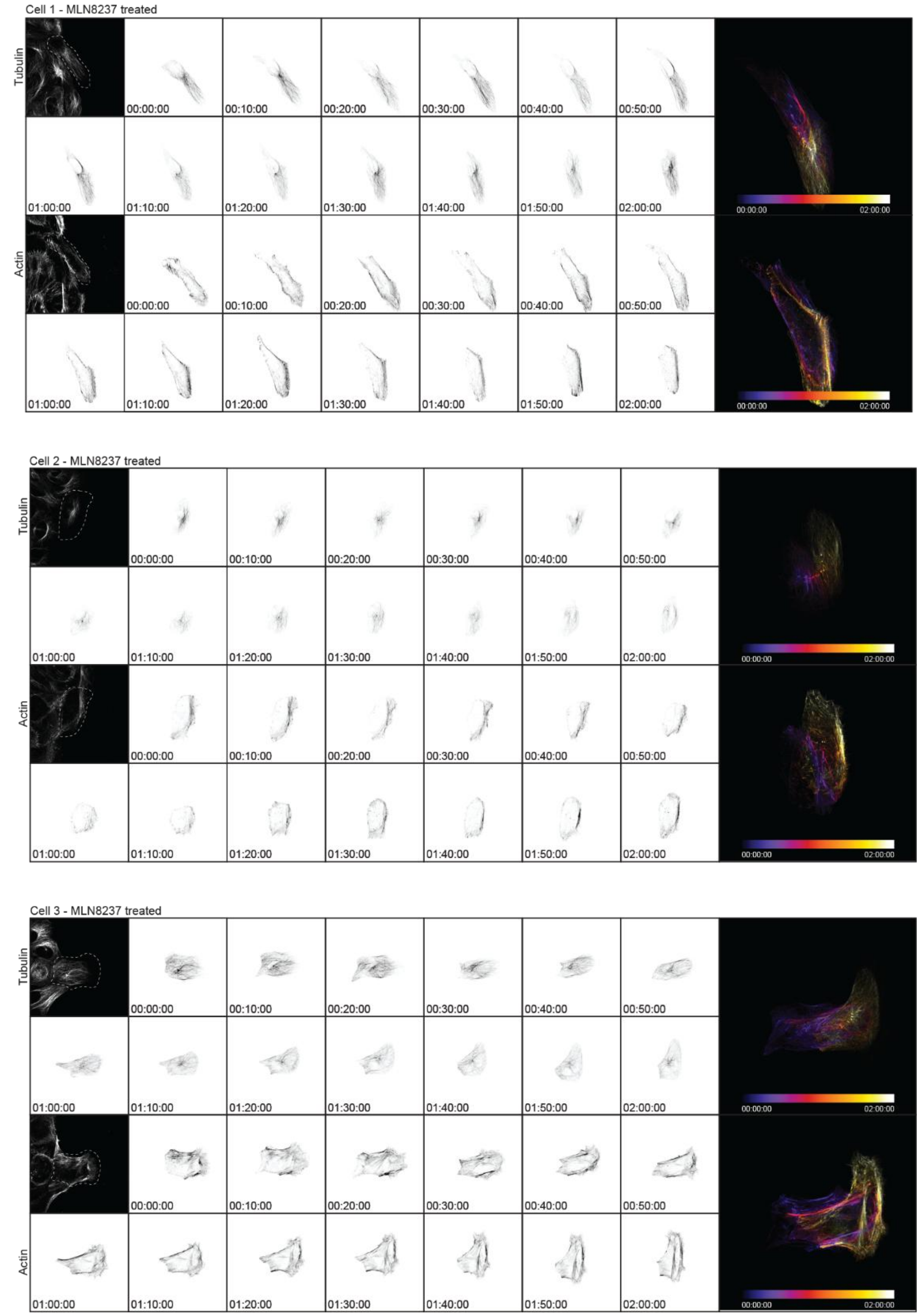


**Supplementary Figure 11. Live-cell imaging of tubulin and actin in MCF10A RFP-TUBA1B leader cells treated with MLN8237.** Live-cell imaging of MCF10A RFP-TUBA1B cells using SPY555-tubulin and SPY650-FastAct. After the addition of MLN8237, cells were monitored for 2 hours at a time interval of 10 minutes. Inverse images of cropped leader cells, then centered around the nucleus (crop is highlighted in dashed lines in the raw images). Time-color coded overlay images of non-centered, cropped leader cells are shown for 2 hours of migration.
